# Improving Split-HaloTag through Computational Protein Engineering

**DOI:** 10.1101/2024.10.13.617931

**Authors:** Jonas Wilhelm, Lennart Nickel, Yin-Hsi Lin, Julien Hiblot, Kai Johnsson

## Abstract

Split-HaloTag can be used to transform transient molecular interactions into permanent marks through chemical labeling, thereby enabling the recording of transient physiological events in individual cells. However, applications of split-HaloTag-based recorders can be limited by slow labeling rates. To address this issue, we have engineered an improved version of cpHaloΔ, the larger fragment of the split-HaloTag system. Using computational techniques, we identified stabilizing point mutations and designed a structured linker connecting the original N and C termini of the circular permutated protein, thereby significantly improving thermostability and activity of cpHaloΔ. These modifications decrease the time and substrate concentrations required for split-HaloTag-based assays and can expand their dynamic range and sensitivity.

## Introduction

Methods that enable the recording of transient cellular events simultaneously across large cellular populations are helpful for investigating complex processes in biological systems^1–3^. For this purpose, we recently developed a split version of the self-labeling protein HaloTag which can be labeled with fluorescent chloroalkane (CA) substrates *in vitro* and *in vivo*^4^. Split-HaloTag was generated by circular permuting HaloTag, followed by the excision of a small helical peptide. The larger protein part (cpHaloΔ) can be efficiently labeled only upon binding of the small peptide (Hpep) (Fig. 1A). An array of different Hpeps (Hpep1-8) with affinities for cpHaloΔ ranging from the nano- to the milli-molar range have been developed, to meet different experimental requirements. By making the reconstitution of functional split-HaloTag dependent on a defined physiological activity, this activity is recorded through an irreversible labeling step that enables *post hoc* analyses. Such recorders can be generated by fusing the split-HaloTag parts to sensing domains, such as calmodulin and its recognition peptide M13, or to G-protein-coupled receptors (GPCRs) and beta-arrestin. Cells that experience elevated calcium levels or signaling events via the tagged GPCR can be selectively and permanently labeled, effectively recording their physiological history^4^. This separation of recording and readout allowed the analysis of large heterogeneous cell populations through information-rich methods such as RNA sequencing, and has enabled brain-wide recordings of neuronal activities in living flies and zebrafish larvae. Furthermore, the availability of fluorescent HaloTag substrates in different colors has facilitated the recording of multiple epochs of neuronal activities in a single individual^4^.

**Figure 1:**
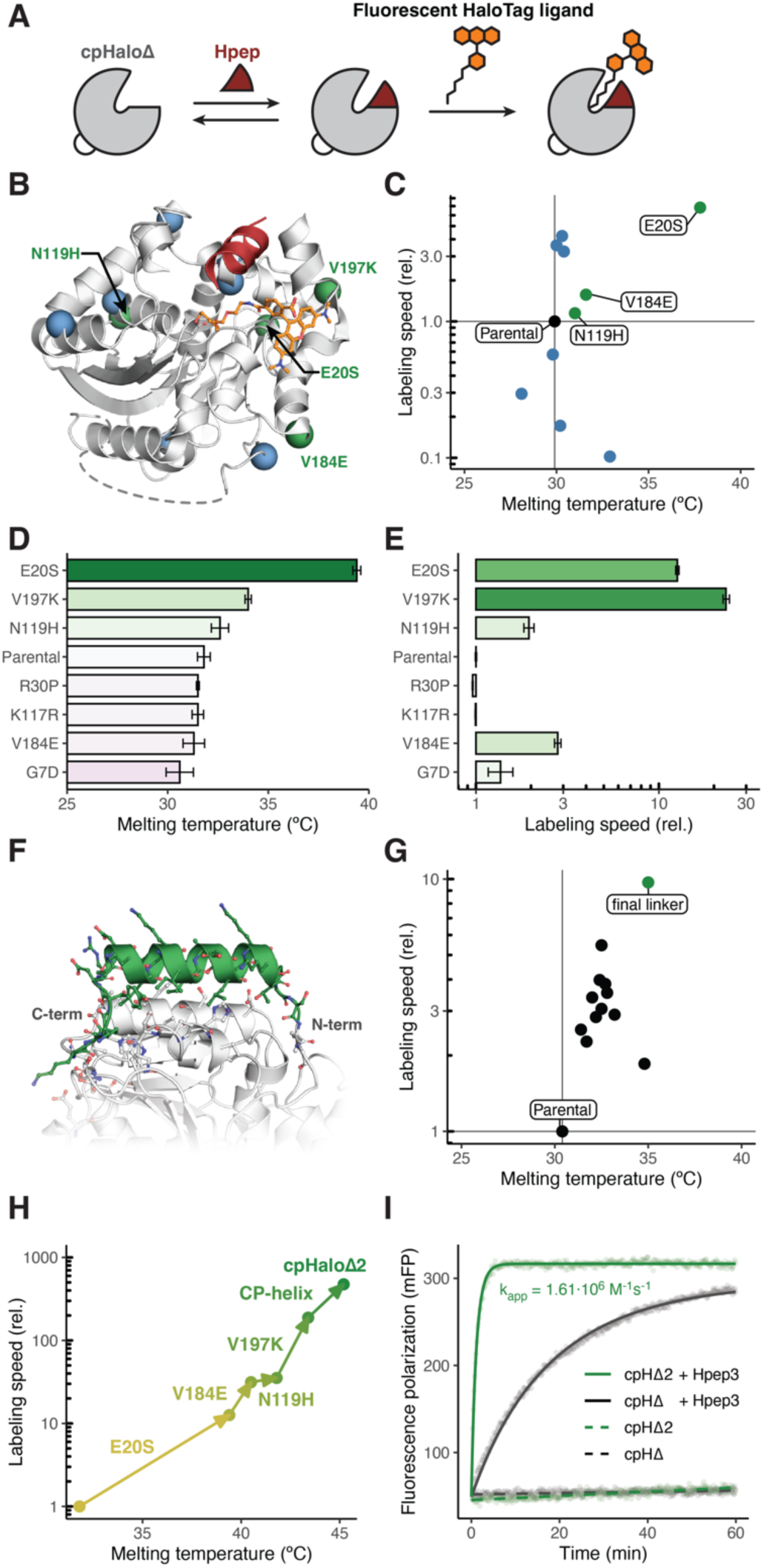
cpHaloΔ2 engineering. (**A**) Schematic of the split-HaloTag system. (**B**) Split-HaloTag model derived from the TMR-CA (orange) labelled HaloTag crystal structure (PDB-ID: 6Y7A^5^). cpHaloΔ is shown in grey, the Hpep in red. Locations of tested mutations from PROSS output are depicted as blue (neutral/negative mutations) and green (beneficial mutations) spheres. (**C**) Melting temperatures and labeling speeds with TMR-CA (20 nM) in presence of Hpep3 (6.25 μM) of tested point mutants (100 nM). Labeling speeds are given relative to the parental cpHaloΔ. (**D**) Melting temperatures of selected cpHaloΔ mutants after removal of purification tags. Error bars represent standard deviations. (**E**) Labeling speeds with TMR-CA (20 nM) in presence of Hpep3 (6.25 μM) of selected cpHaloΔ mutants (100 nM) after removal of purification tags. Labeling speeds are given relative to the parental cpHaloΔ. Error bars represent 95% confidence intervals. (**F**) Structure of cpHaloΔ (grey) with designed helical linker (green) connecting the original termini. (**G**) Melting temperatures and labeling speeds with TMR-CA (20 nM) in presence of Hpep3 (6.25 μM) of cpHaloΔ variants (100 nM) with different designed linkers. Labeling speeds are given relative to the parental cpHaloΔ (**H**) Melting temperatures and labeling speeds with TMR-CA (20 nM) in presence of Hpep3 (6.25 μM) of cpHaloΔ variants (100 nM) combining beneficial mutations and linkers. Labeling speeds are given relative to the parental cpHaloΔ, purification tags of protein variants were removed. (**I**) Labeling kinetics of cpHaloΔ and cpHaloΔ2 (10 nM) with TMR-CA (2 nM) in presence or absence of Hpep3 at saturating concentrations (1.5 mM) followed by fluorescence polarization. A second-order reaction model was fitted to the data to estimate apparent second-order rate constants (k_app_).

Despite the successful applications of split-HaloTag, a considerable limitation of the approach can be the relatively long labeling time required to achieve well-detectable signals, particularly when effective substrate concentrations are limited, such as in animal models^4^. This reduces the achievable temporal resolution, as labeling periods ranging from several minutes to hours might be required to record a detectable signal. While HaloTag labeling is very fast and efficient, reaching apparent second-order rate constants above 10^7^ M^-1^s^-1^ for certain substrates^5^, labeling of the fully complemented split-HaloTag is significantly slower at about 10^4^ M^-1^s^-1^ ^4^. This suggests that the speed and efficiency of split-HaloTag labeling could be enhanced, which would facilitate shorter labeling times and lower substrate concentrations, ultimately improving the temporal resolution of split-HaloTag-based assays.

## Results

One reason for the lower activity of split-HaloTag compared to the native HaloTag could be a decrease in thermostability of the large fragment (cpHaloΔ). Such destabilization could be expected as many split enzymes feature decreased stability due to exposed hydrophobic residues after splitting^6^. Indeed, we measured a melting temperature of 31.8 °C for cpHaloΔ which is far below the melting temperature of the native HaloTag at 62.0 °C. This indicates that at the physiological temperature of 37 °C a significant fraction of the protein might be in an unfolded state, not available for complementation with the Hpep and subsequent labeling. We hypothesized that stabilizing cpHaloΔ could increase the fraction of active protein, and hence benefit experiments by increasing labeling rates. This improvement would enable shorter assay durations and reduce the amount of fluorophore required at the site of action. In order to stabilize the protein, we used the PROSS^7^ workflow to identify potentially beneficial point mutations using the TMR (tetramethyl rhodamine) labeled HaloTag crystal structure as input (PDB-ID: 6Y7A^5^). PROSS leverages evolutionary information and physics-based energy calculations through Rosetta to find mutations that may enhance stability. From the PROSS output, we selected twelve mutations, excluding those near the active site, the Hpep binding site, the substrate binding channel, and the rhodamine binding site on the protein surface (Fig. 1B).

Ten out of the twelve single point mutants expressed well in *E.coli* and could be purified, while one mutant (V197K, according to HaloTag numbering) could not be cloned and another one (I135F) did not express. Mutants were screened for thermostabilities and labeling rates in the presence of a non-saturating concentration of Hpep3. All labeling experiments were conducted with a 20-minute pre-incubation time at 37 °C to mimic physiological conditions. Eight mutants showed increased melting temperatures among which six exhibited faster labeling rates at 37 °C (Fig. 1C, S1, S2). Two of the stabilizing mutations had detrimental effects on labeling rates.

The largest effect was found for mutation E20S, which led to an increase in melting temperature of 7.9 °C and a 6.8-fold increase in labeling rate during screening. To simplify the screening procedure, assays were performed with N-terminally His-tagged proteins. However, the His-tag is directly adjacent to the Hpep binding site of cpHaloΔ, potentially impacting Hpep binding and interfering with the labeling reaction. To rule out such effects, the six mutants that showed beneficial properties in the initial screen were tested without purification tags, which were removed by tobacco etch virus protease (TEVp) digestion. At this stage, we included the previously missing mutant V197K. For three mutants (E20S, N119H, and the newly included V197K) the positive effect on melting temperature and labeling rate was confirmed (Fig. 1D, E). Notably, the V197K mutant featured a 23.3-fold increase in labeling rate at 37 °C, while E20S increased melting temperature by 7.6 °C. While having no clear effect on thermostability, the V184E mutation led to a 2.8-fold increase in labeling rate and was hence also considered as beneficial.

As an additional strategy to improve cpHaloΔ stability and activity, we designed a new linker connecting the original N- and C-termini of the circularly permuted HaloTag. We reasoned that replacing the flexible (GGS/T)_5_ linker with a well-structured alpha-helical linker, which interacts with both the N- and C-terminal parts of cpHaloΔ, could assist in optimally aligning the parts and rigidifying the intramolecular complex. We used the RosettaRemodel^8^ application to design structures and sequences that would fulfill these criteria. To accommodate the formation of an alpha helix, we increased the linker length from 15 to 22 or 23 residues. In total 20,000 structures were generated and scored by Rosetta (Fig. 1F). The top six scoring designs for each linker length were tested experimentally. All tested linkers positively impacted thermostability and labeling speed at 37°C, with one linker standing out, resulting in a 4.6°C increase in melting temperature and a 9.7-fold faster labeling rate at 37°C (Fig. 1G, S3, S4). Notably, this linker had the best Rosetta score of all 23 residue linkers, and its design closely matched its AlphaFold 3^9^ predicted structure (RMSD 0.530 Å, Fig. S5).

We then combined the identified beneficial mutations (E20S, N116H, V184E, V197K) and the improved circular permutation linker. The resulting protein, termed cpHaloΔ2, features a melting temperature of 45.2 °C (an increase of 15 °C) and a 475-fold higher labeling speed in the presence of Hpep3 (Fig. 1H). However, the reported labeling rates were determined at a non-saturating concentration of Hpep3, hence the observed increase in labeling speed may result from both, enhanced labeling activity of the complemented species and increased affinity of cpHaloΔ2 for the Hpep3. At saturating Hpep3 concentration, we measured an apparent second-order rate constant of 1.61 ⋅ 10^6^ M^-1^s^-1^ for cpHaloΔ2, which is 16 times higher than that of the original cpHaloΔ under the same conditions (Fig. 1I). This suggests that in addition to higher activity of the complemented protein, the introduced modifications may also increase the affinity of cpHaloΔ2 for the Hpep. To estimate affinities, we determined EC_50_ values of Hpep variants 1-8 for the cpHaloΔ2 labeling reaction (Fig. 2A, S6). In comparison to the original cpHaloΔ, a median decrease in EC_50_ by a factor of 3.5 was observed (Fig. 2B). Thus, the improved cpHaloΔ2 still covers a wide range of affinities for Hpep variants 1-8, ranging from 43 nM to 2.4 mM.

**Figure 2:**
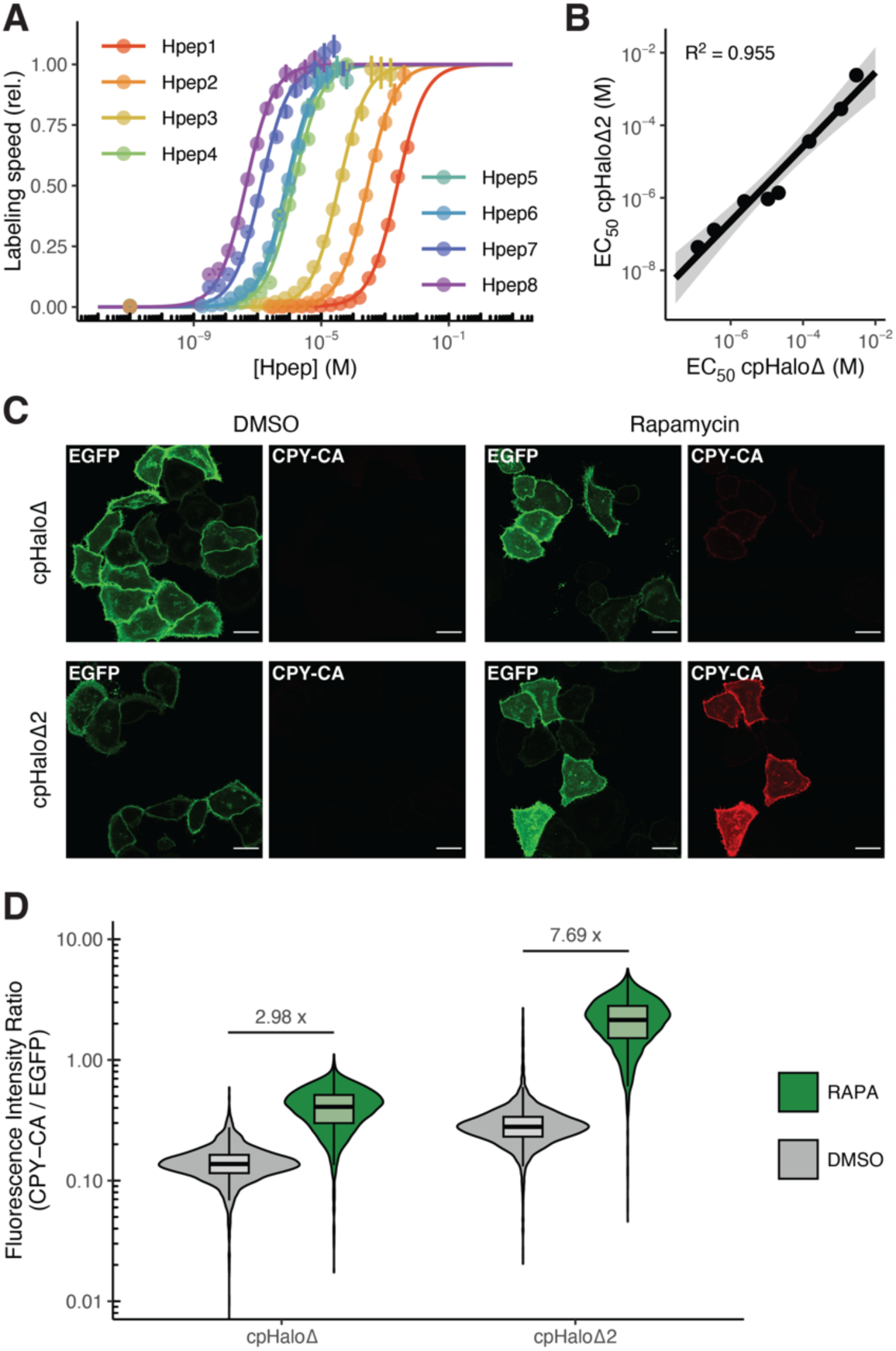
cpHaloΔ2 characterization. (**A**) Relative labeling rates of cpHaloΔ2 (10 nM) with TMR-CA (2 nM) at different concentrations of Hpep variants 1-8. A sigmoidal model was fitted to the data to estimate EC_50_ values. EC_50_ values are ranging from 43 nM to 2.4 mM. Error bars represent 95% confidence intervals. (**B**) Comparison of EC_50_ values between cpHaloΔ and cpHaloΔ2. A linear model was fitted to the log transformed data. EC_50_ values for cpHaloΔ2 show a median decrease by a factor of 3.5. (**C**) Confocal fluorescence micrographs of HeLa cells co-expressing Lyn11-EGFP-cpHaloΔ/cpHaloΔ2-(GGS)_9_- FKBP and Hpep3-(GGS)_3_-FRB-mScarlet. Labeling with CPY-CA (10 nM, 30 min) is observed only for the cpHaloΔ2 construct in the presence of rapamycin (100 nM). Scale bars are 25 μm. (**D**) Flow cytometry analysis of HeLa cells co-expressing Lyn11-EGFP-cpHaloΔ/cpHaloΔ2-(GGS)_9_-FKBP and Hpep3-(GGS)_3_-FRB-mScarlet labeled with CPY-CA (10 nM, 30 min) in the presence or absence of rapamycin (100 nM). The cpHaloΔ2 construct shows 5.3-fold stronger labeling in presence of rapamycin compared to cpHaloΔ. cpHaloΔ2 also displays a larger difference in labeling with or without rapamycin.

Next, we investigated the effect of cpHaloΔ stabilization on the background labeling in absence of Hpep. We measured an apparent second-order rate constant of 3.26 M^-1^s^-1^, which is a 15.5-fold increase in comparison to the original cpHaloΔ. Hence, background labeling and labeling of the complemented protein increased by similar factors, leaving the large dynamic range of the system unchanged.

To demonstrate the performance of cpHaloΔ2 in live cells, we fused the split-HaloTag fragments to FKBP and FRB, which interact selectively upon the addition of the small molecule dimerizer rapamycin. The split parts were fused to the N-termini of FKBP and FRB, which are separated by 49 Å in the complex, representing a realistic use case, as split-HaloTag fusion partners in specific assays may often have non-adjacent termini. We co-expressed Hpep3-(GGS)_3_-FRB-mScarlet and either Lyn11-EGFP-cpHaloΔ-(GGS)_9_-FKBP or Lyn11-EGFP-cpHaloΔ2-(GGS)_9_-FKBP in HeLa cells and labeled them in presence of rapamycin at a low concentration of CPY-CA (carbopyronine-CA, 10 nM) fluorescent substrate. After 30 minutes, strong labeling at the plasma membrane was observed only with cpHaloΔ2, while the original cpHaloΔ showed only minimal labeling under these conditions (Fig. 2C). Both versions did not exhibit significant labeling in the absence of rapamycin. To further quantify their labeling, we subjected cells treated as described above to flow cytometry analysis. In the presence of rapamycin, cpHaloΔ2 exhibited a 5.3-fold higher median labeling than the original cpHaloΔ. Additionally, cpHaloΔ2 showed a 2.6-fold greater difference in labeling intensity with or without rapamycin (Fig. 2D). These experiments demonstrate that the improvements in stability and labeling speed translate to higher signal intensities and a more sensitive assay in living cells.

## Discussion

We have developed an improved version of the split-HaloTag by increasing the stability and activity of the cpHaloΔ fragment through computational protein engineering. The improved split system reacts more efficiently with fluorescent HaloTag substrates both *in vitro* and in living cells, reducing required incubation times and substrate concentrations. This addresses a critical need, as the application of split-HaloTag-based molecular recorders has been limited by prolonged incubation times in animal models. We anticipate that such experiments will benefit from the improved split-HaloTag system, effectively increasing the temporal resolution of the approach. This could broaden the scope of possible experiments and particularly improve the feasibility of performing multiple recordings in a single individual using different colors of HaloTag ligands.

We expect that the improved split-HaloTag will not only improve existing recorders, but also facilitate the development of new molecular recorders for various biological processes. Specifically, the characterization of labeling speeds, background reactivity and split protein affinity, along with the availability of Hpep sequences with affinities across the nanomolar to millimolar range, should aid in the design of novel split-HaloTag-based tools. Furthermore, the described computational method for improving circularly permuted proteins by identifying stabilizing point mutations and designing circular permutation linkers with defined secondary structures could serve as a blueprint for enhancing other circularly permuted proteins, which are widely used in biosensors and as reporters^10^.

## Materials and Methods

### General information

If not specified otherwise, reagents were obtained from Merck KGaA and used without further purification. All microplate reader measurements were performed with a Spark 20M instrument (Tecan Group Ltd.). Fluorescent HaloTag ligands (TMR-CA and CPY-CA) were prepared in house according to published procedures^11,12^.

### Identification of point mutations with PROSS

PROSS^7^ was run via the web interface (https://pross.weizmann.ac.il) with default settings.

### Design of circular permutation linkers

The cpHaloΔ structure was first minimized using the Rosetta FastRelax^13^ application with the following flags:

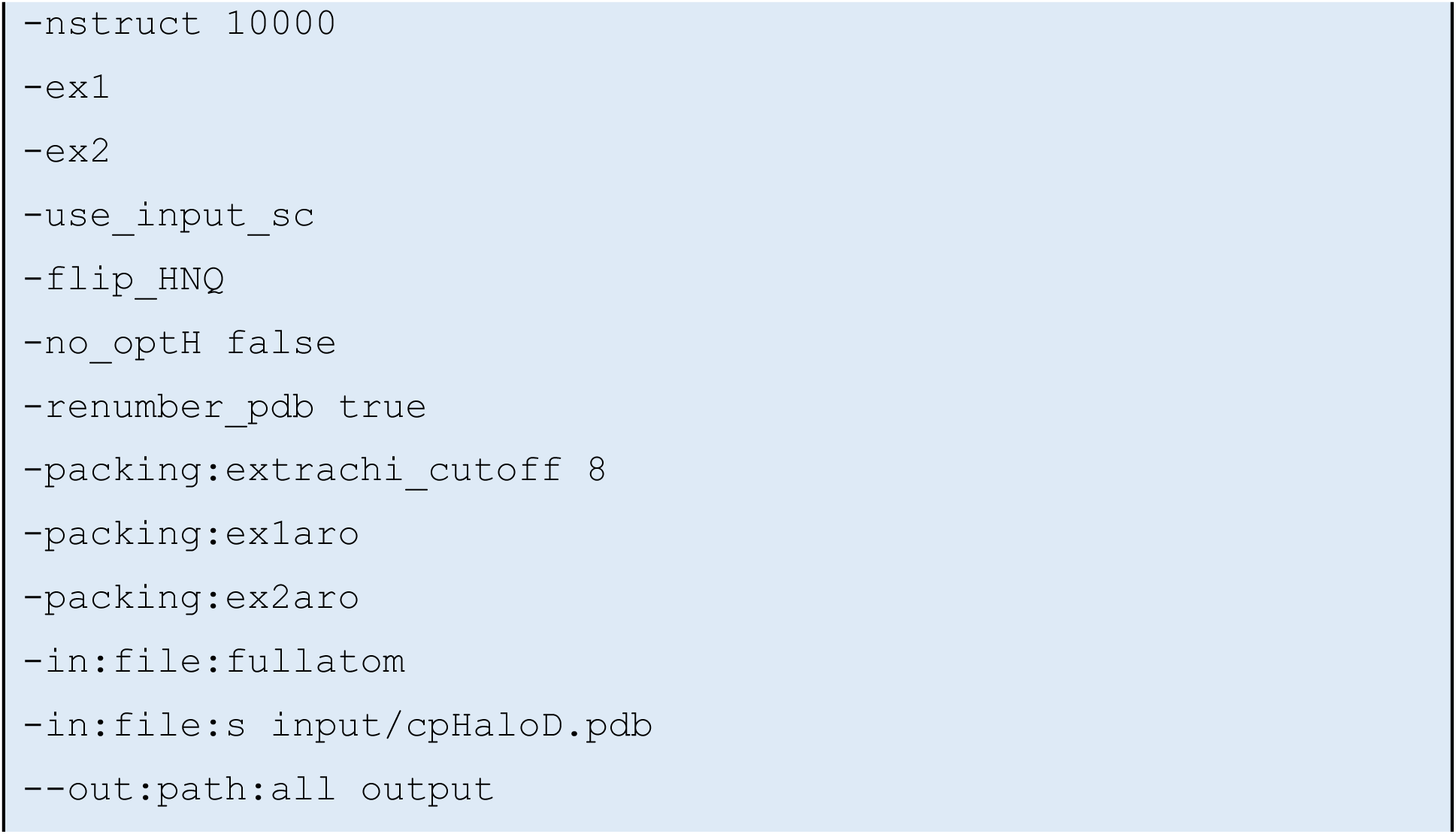

The top-scoring structure was used as input for designing circular permutation linkers using the RosettaRemodel^8^ application. Blueprint files for 22 and 23 amino acid helical linkers were set up, allowing any amino acids except cysteine in the designed regions. 22 amino acid linker blueprint:

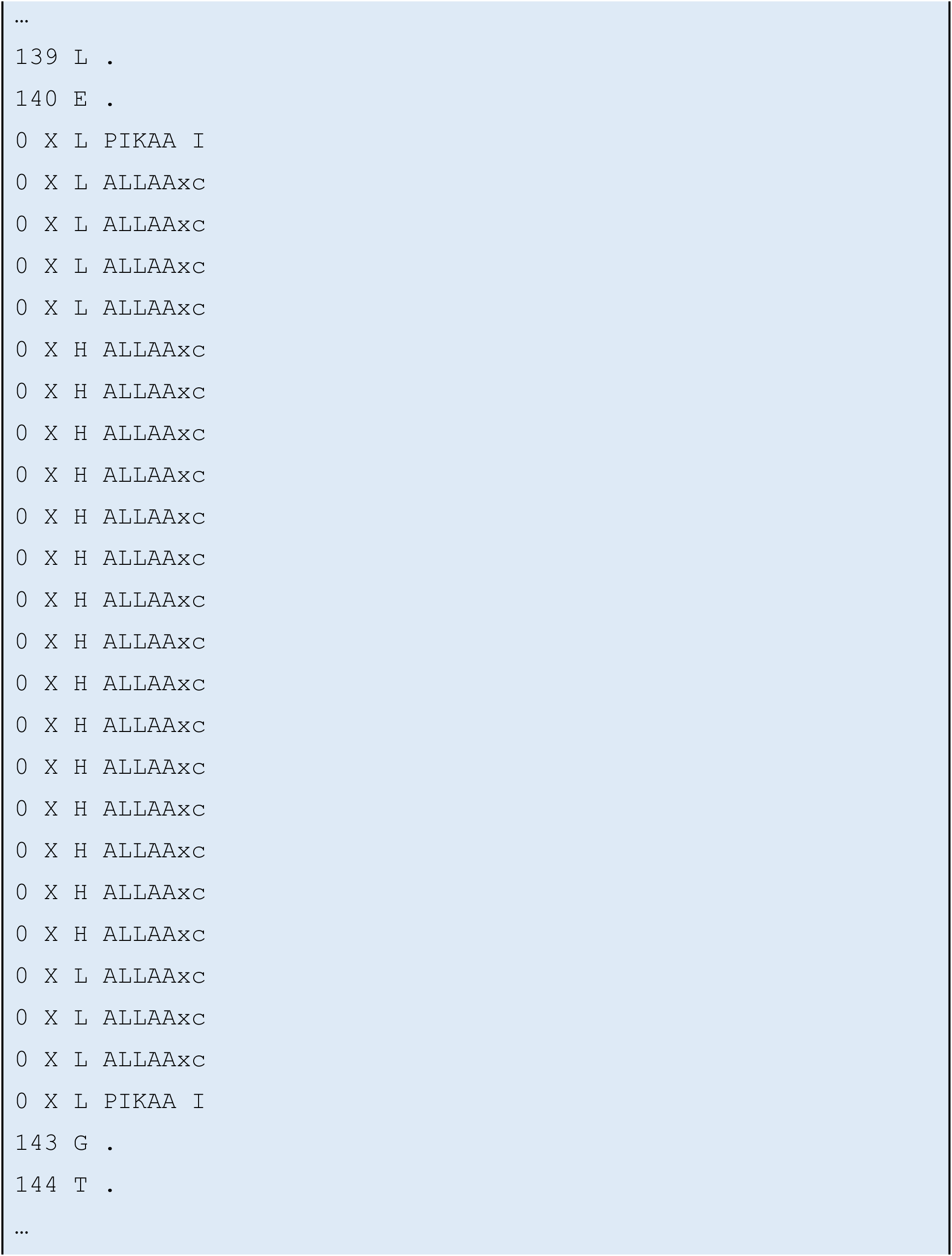

23 amino acid linker blueprint:

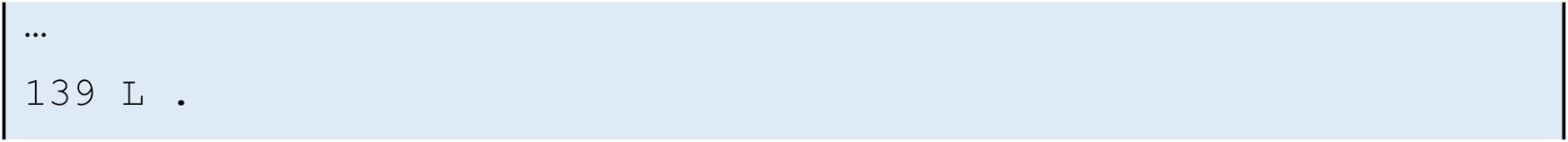
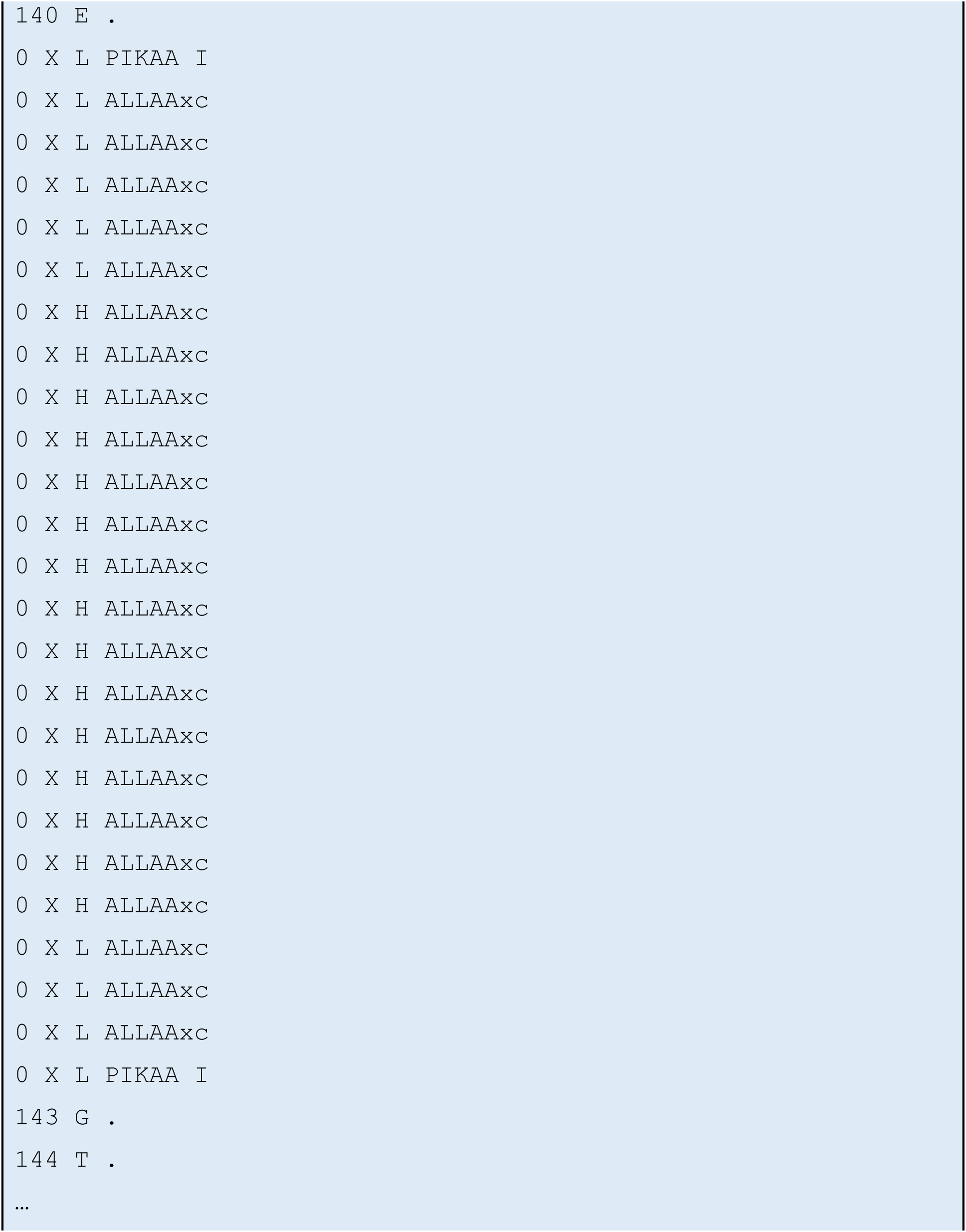

RosettaRemodel was run with the following flags:

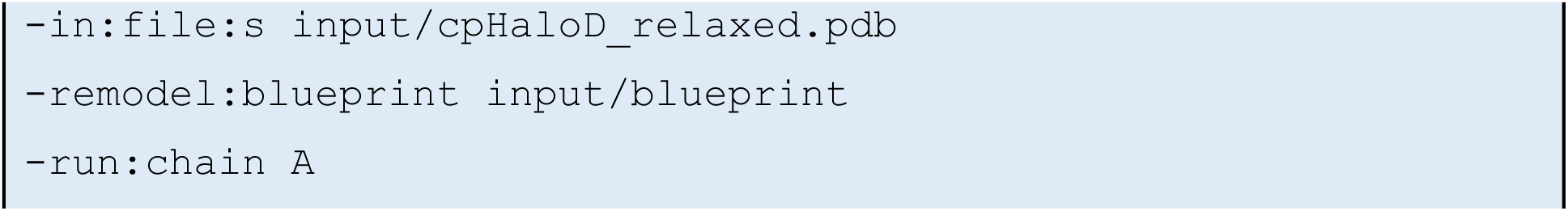
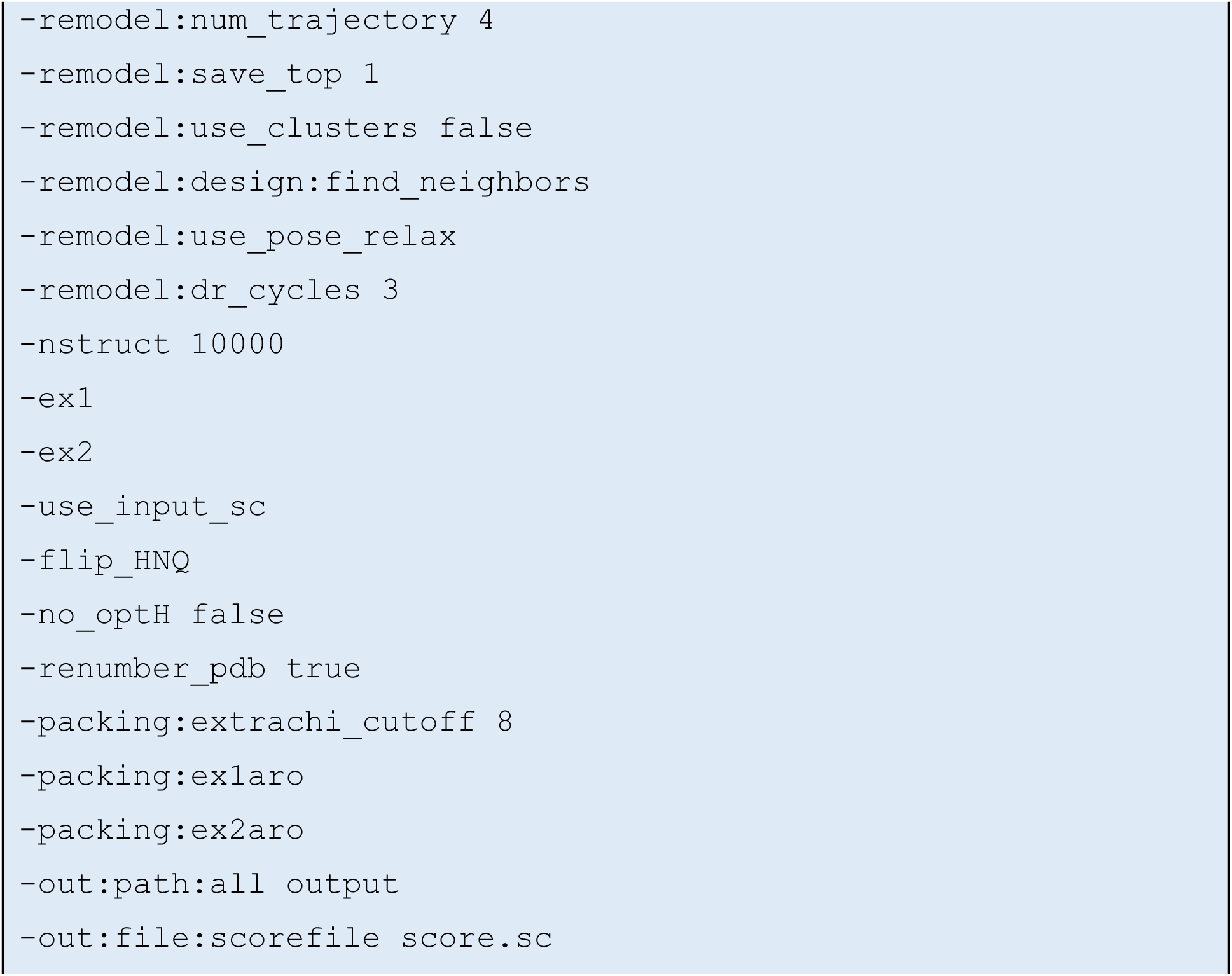

The top six scoring sequences for both, the 22 and 23 amino acid linkers were tested experimentally. The input and top-scoring output structures can be found in the supplementary material (‘Structures.zip’).

### Molecular cloning

Plasmids for bacterial expression were prepared with the Q5 site-directed mutagenesis kit (New England Biolabs Inc.) according to the manufacturers’ protocol. Sequences were cloned in a modified pET-51b(+) vector with an N-terminal His_10_-tag followed by a TEVp cleavage site and no C-terminal tags. Plasmids for mammalian expression were prepared using Gibson assembly^14^. Sequences were cloned in the pcDNA™5/FRT (Thermo Fisher Scientific Inc.) vector. All constructs were confirmed via sanger sequencing.

### Protein expression and purification

Proteins were expressed in *E.coli* BL21 (DE3) (Merck KGaA). Individual clones were grown in lysogenic broth (LB) at 37 °C, 220 rpm until an optical density at 600 nm of 0.6-0.8 was reached. Cultures were cooled (10 min, 8 °C) and transgene expression was initiated by addition of 0.5 mM isopropyl-β-D-thiogalactopyranoside (IPTG). Cultures were grown at 16 °C, 220 rpm for 16-20 h. Cells were harvested by centrifugation (5000 g, 10 min, 4 °C), resuspended in 30 mL lysis buffer (50 mM KH_2_PO_4_, 150 mM NaCl, 5 mM imidazole, 1 mM phenylmethylsulfonyl fluoride (PMSF), 0.25 mg/mL lysozyme, pH 8.0) and lysed by sonication on wet ice. Lysates were cleared by centrifugation (70 000 g, 20 min, 10 °C) and proteins were purified via immobilized metal affinity chromatography (IMAC) using a HisTrap FF crude column on an ÄktaPure FPLC system (Cytiva Europe GmbH). Columns were washed with 50 mM KH_2_PO_4_, 300 mM NaCl, 10 mM imidazole, pH 7.5 and proteins were eluted in 50 mM KH_2_PO_4_, 300 mM NaCl, 500 mM imidazole, pH 7.5. Buffer was exchanged on a HiPrep 26/10 desalting column (Cytiva Europe GmbH) to 50 mM HEPES, 50 mM NaCl, pH 7.3. Proteins were concentrated using Amicon Ultra-15 centrifugal filter devices (Merck) with a molecular weight cut-off of 30 kDa to a final concentration of 100-500 μM. Correct size and purity of proteins were assessed by SDS-PAGE and liquid chromatography mass spectrometry (LCMS) analysis. Mass spectrometry measurements were performed on a Bruker maXis II ETD connected to a Shimadzu Nexera X2 HPLC.

### Removal of purification tags by TEVp cleavage

Cleavage with tobacco etch virus protease (TEVp) was performed in TEVp buffer (25 mM KH_2_PO_4_, 200 mM NaCl, 10% glycerol, 5 mM 2-mercaptoethanol, pH 8.0) overnight at 4 °C. Samples were filtered (0.22 μM) and purified by IMAC, collecting the flow-through. Proteins were concentrated as described above and further purified by size exclusion chromatography with a HiLoad 26/600 Superdex 75 pg column on an ÄktaPure FPLC system (Cytiva Europe GmbH) in 50 mM HEPES, 50 mM NaCl, pH 7.3. Proteins were concentrated and correct size and purity were assessed as described above.

### Protein thermostability measurements

Melting temperatures were measured in in technical duplicates at 20 μM protein in HEPES buffer (50 mM HEPES, 50 mM NaCl, pH 7.3) on a Prometheus NT 48 nanoscale differential scanning fluorimeter (NanoTemper Technologies GmbH) over a temperature range of 20-95 °C with a heating rate of 1 °C min^−1^ by recording changes in the ratio of tryptophane fluorescence at 350 nm and 330 nm. Melting temperatures were determined as the inflection point, *i.e.* maximum of the first derivative, of the fluorescence intensity ratio.

### Split-HaloTag labeling kinetics

Split-HaloTag labeling kinetics with TMR-CA were measured in FP buffer (50 mM HEPES, 50 mM NaCl, 0.1 g/L bovine serum albumin (BSA), pH 7.3) by recording fluorescence polarization over time in a plate reader. Assays were performed at 37 °C in black, non-binding, flat-bottom 384-well plates with a final volume of 40 μL. Plates with all reagents except fluorescent substrate were equilibrated at 37 °C for 20 min and reactions were started by the addition of TMR-CA using an electronic 384-channel pipettor (VIAFLO, Integra Biosciences GmbH). A humidity cassette (Tecan Group Ltd.) was used to reduce evaporation during the measurement. Screenings for circular permutation linkers were performed once, point mutations and combinations of mutations were screened in technical duplicates. Measurements for determining EC_50_ values for different Hpeps and for direct comparison between cpHaloΔ and cpHaloΔ2 were performed in technical triplicates.

A second-order reaction model (equation 1) was fitted to the data to determine apparent second-order rate constants (k_app_).

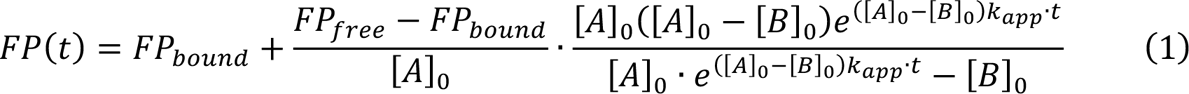

With:

t: time

FP(t): FP at time t

FP_free_: FP of the free dye

FP_bound_: FP of the bound dye [A]_0_: dye concentration at t = 0

[B]_0_: protein concentration at t = 0

k_app_: apparent second-order rate constant

In cases where the reaction did not reach a plateau a linear model was fitted to determine initial slopes (s_t=0_). k_app_ values were estimated from initial slopes using the derivative of equation X at t = 0 (equation 2).

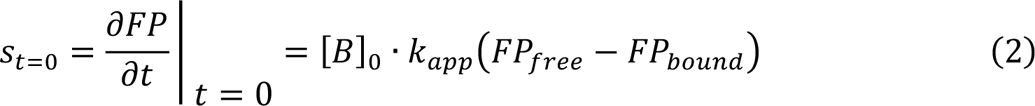

With:

t: time

s_t=0_: initial slope

FP_free_: FP of the free dye

FP_bound_: FP of the bound dye

[B]_0_: protein concentration at t = 0

k_app_: apparent second-order rate constant

Confidence intervals of fitted parameters were estimated with the Monte Carlo method^15^.

### Half-maximal effective concentrations (EC_50_) of Hpeps

cpHaloΔ labeling kinetics were measured at a range of Hpep concentrations as described above to determine apparent reaction rates. Rates were plotted against Hpep concentration and a sigmoidal model (equation 3) was fitted to the data to determine EC_50_ values.

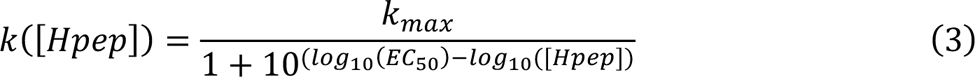

With:

k: apparent second-order rate constant

k_max_: maximal apparent second-order rate constant (at saturating [Hpep])

EC_50_: half-maximal effective concentration

[Hpep]: Hpep concentration

Confidence intervals of fitted parameters were estimated with the Monte Carlo method^15^.

Background labeling kinetics of cpHaloΔ

Labeling kinetics of cpHaloΔ with TMR-CA in absence of the Hpep were measured in FP buffer (50 mM HEPES, 50 mM NaCl, 0.1 g/L BSA, pH 7.3) by recording fluorescence polarization over time in a plate reader. Assays were performed at 37 °C in black, non-binding, flat-bottom 384-well plates with a final volume of 40 μL in technical triplicates. Plates were equilibrated at 37 °C for 20 min and reactions were started by the addition of TMR-CA using an electronic 384-channel pipettor (VIAFLO, Integra Biosciences GmbH). A humidity cassette (Tecan Group Ltd.) was used to reduce evaporation during the measurement. A two-step reaction model (equations 4, 5) was fitted to the data globally using the DynaFit software^16^ to determine 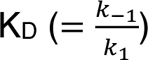, k_2_ and k_app_ (= *k*_>_ ⋅ *k*_2_/(*k*_?_ + *k*_−1_)) of the background labeling reaction. Confidence intervals of fitted parameters were estimated with the Monte Carlo method^15^.

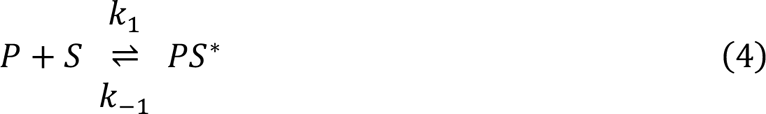

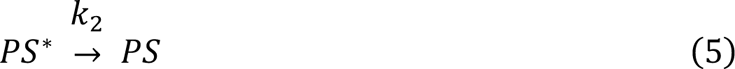

### Labeling of rapamycin-dependent FKPB-FRB interactions in HeLa cells

HeLa cells were cultured in DMEM + GlutaMax (Gibco, Thermo Fisher Scientific Inc.) medium supplemented with glucose (4.5 g/L), pyruvate (110 mg/L) and 10% FBS at 37 °C in a humidified incubator with 5% CO_2_ atmosphere. Cells were transiently transfected with split-HaloTag-FKBP/FRB co-expression constructs using Lipofectamine 3000 (Thermo Fisher Scientific Inc.) according to the manufacturer’s protocol. After 8 h incubation, the medium was replaced and cells were grown for additional 24 h. Cells were treated with 10 nM CPY-CA in the presence or absence of 100 nM rapamycin for 30 min. Cells were washed 3 times with fresh medium containing 1 μM HaloTag protein to scavenge the substrate before analysis via flow cytometry or confocal fluorescence microscopy.

For confocal microscopy, cells were washed with PBS, fixed with 4% PFA in PBS for 15 min and washed again with PBS. Confocal microscopy was performed on a Stellaris 5 inverted microscope (Leica Microsystems GmbH) equipped with a white line laser and hybrid photodetectors at 37 °C. A 40x/1.10 water immersion objective was used at to image a single plane at 1024×1024 pixels (194×194 μm) resolution (1.5 x zoom, 600 Hz scan speed, 68.8 μm pinhole, 6-fold line average, mEGFP 488 nm ex., 494 nm – 556 nm em., CPY 610 nm ex., 620 nm – 750 nm em.). Brightness of images was adjusted equally for all recorded images using FIJI.

For flow cytometry cells were detached using TrypLE™-Express (Gibco, Thermo Fisher Scientific Inc.) for 5 min at 37 °C. Detached cells were suspended in PBS containing 2% FBS. Cells were analyzed with a BD Fortessa X-20 flow cytometer (mEGFP: 488 nm excitation, 530/30 nm emission, CPY: 648 nm excitation, 660/20 nm emission). Live (SSC-A/FSC-A), single (SSC-H/SSC-A) and mEGFP positive cells were gated and fluorescence intensity ratios (CPY-CA/mEGFP) were calculated.

## Supporting information

Structures.zip

## Acknowledgements

We thank Dr. M. Tarnawski for protein melting temperature measurements. We thank A. Bergner and B. Réssy for reagents and materials. We thank Dr. Sebastian Fabritz with the mass spectrometry core facility of the Max-Planck-Institute for medical research for mass spectrometry measurements. J.W. acknowledges support from the Heidelberg Biosciences International Graduate School (HBIGS).

This work was supported by the Max Planck Society (J.W., L.N., Y.H.L., J.H., K.J.); Ecole Polytechnique Federale de Lausanne (EPFL) (K.J.), the Max Planck School Matter to Life (J.W.) and the EPFL Doctoral Program in Biotechnology and Bioengineering (EDBB) (Y.H.L).

## Supplementary material content

PDF file ‘Supplementary_Material.pdf’

Figures S1 to S8

Tables S1 to S3

Protein sequences

ZIP archive ‘Structures.zip’

Input structure for RosettaRemodel based linker design

Top-scoring output structures from RosettaRemodel based linker design

AlphaFold 3 model of cpHaloΔ2

