## Supplementary_Material.pdf for "Improving Split-HaloTag through Computational Protein Engineering"

**This PDF file includes:**

Supplementary figures S1 to S8

Supplementary tables S1 to S3

Protein sequences

### Supplementary Figures

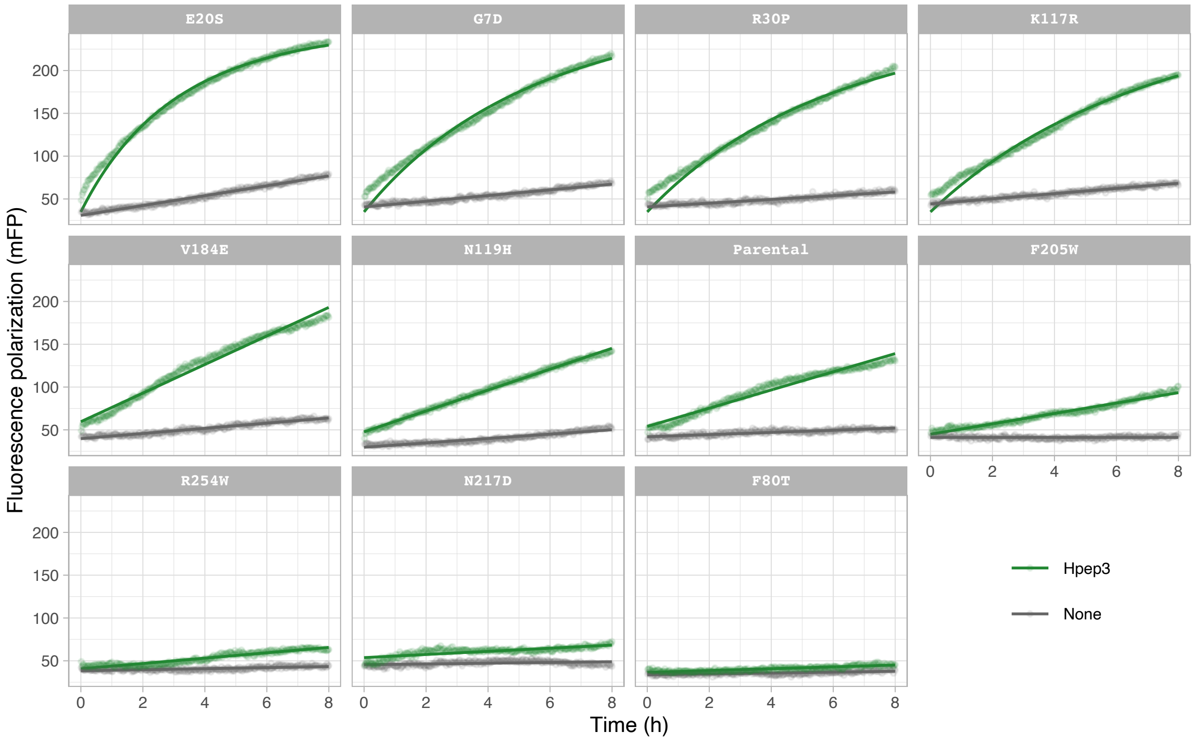


**Fig. S1. Labeling kinetics of cpHaloΔ point mutants.**

Fluorescence polarization labeling kinetics of cpHaloΔ point mutants (100 nM) with TMR-CA (20 nM) in presence or absence of Hpep3 (6.25 µM). Second-order reaction models or linear models (if reactions did not plateau) were fitted to the data to determine labeling rates.

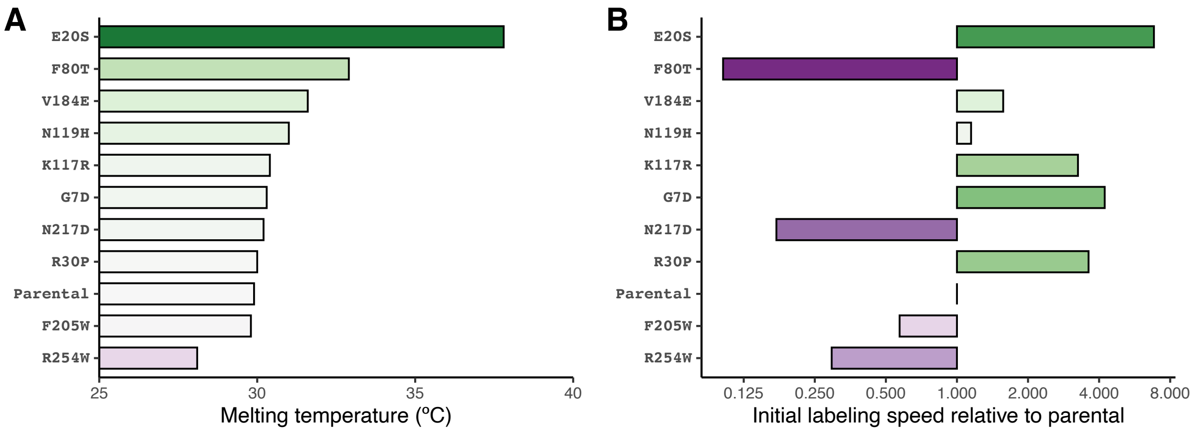


**Fig. S2. Melting temperatures and relative labeling speeds of screened cpHaloΔ point mutants.**

Melting temperatures were determined by nanoDSF, labeling rates via fluorescence polarization (see Fig. S1) at non-saturating Hpep concentrations.

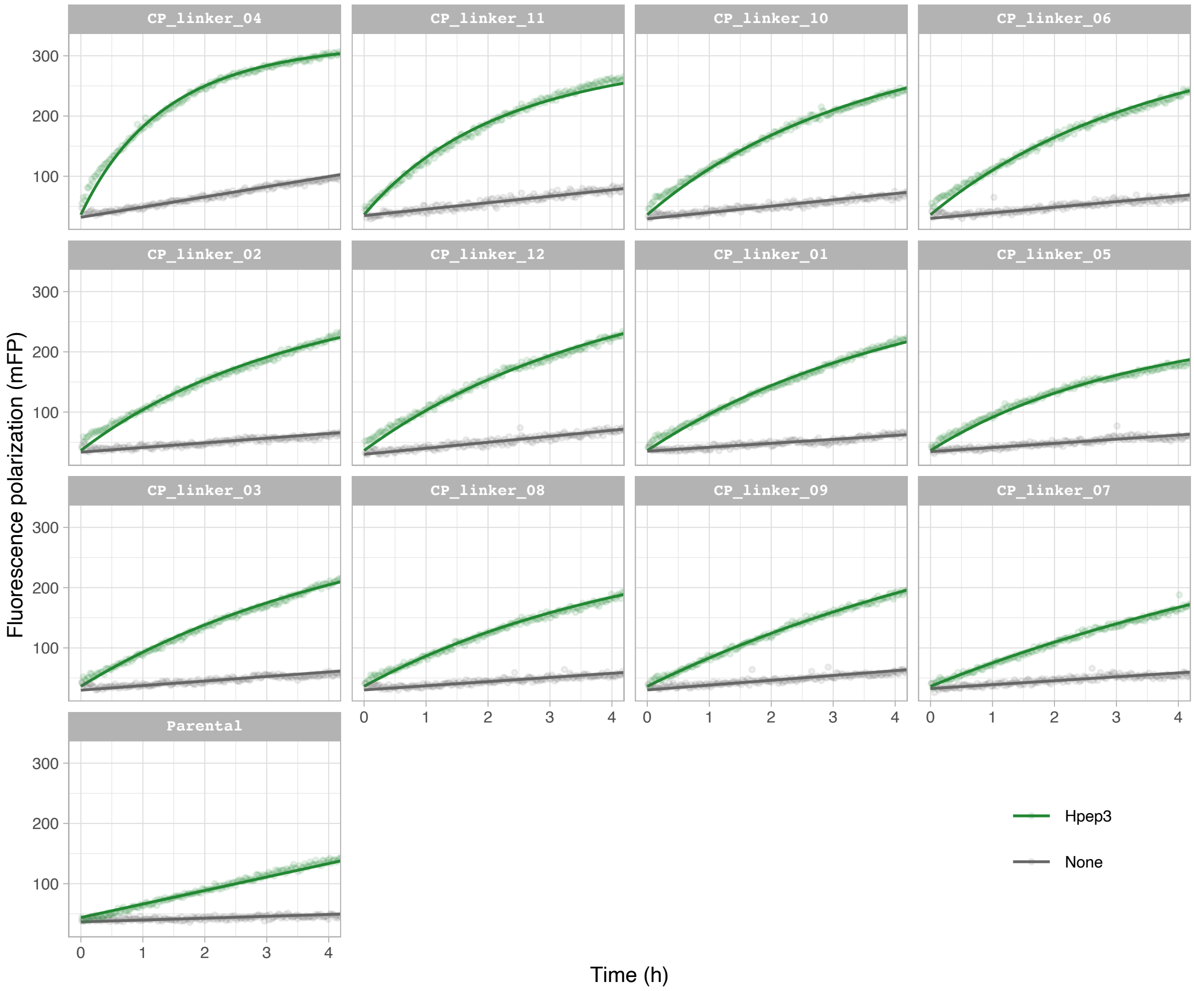


**Fig. S3. Labeling kinetics of cpHaloΔ variants with designed circular permutation linkers.**

Fluorescence polarization labeling kinetics of cpHaloΔ variants (100 nM) with TMR-CA (20 nM) in presence or absence of Hpep3 (6.25 µM). Second-order reaction models or linear models (if reactions did not plateau) were fitted to the data to determine labeling rates.

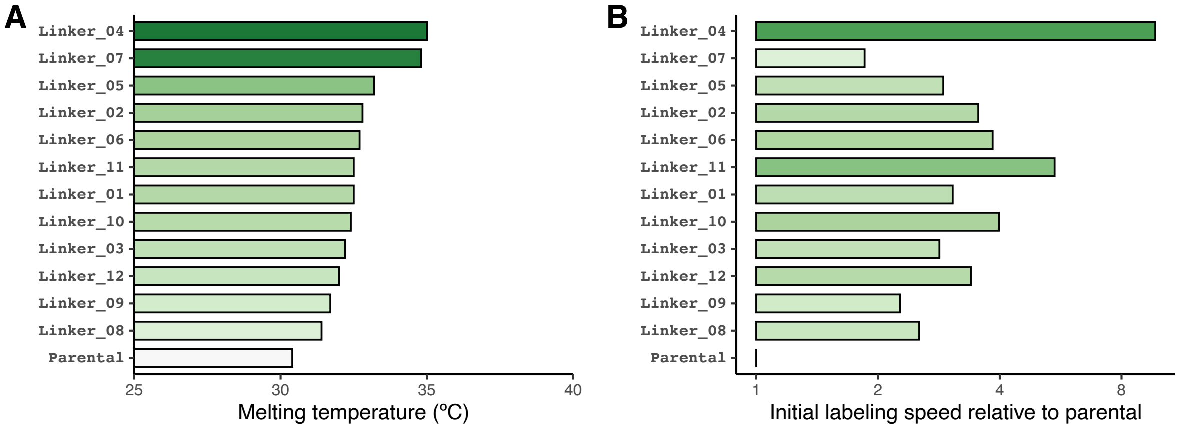


**Fig. S4. Melting temperatures and relative labeling speeds of screened cpHaloΔ variants with designed circular permutation linkers.**

Melting temperatures were determined by nanoDSF, labeling rates via fluorescence polarization (see Fig. S3) at non-saturating Hpep concentrations.

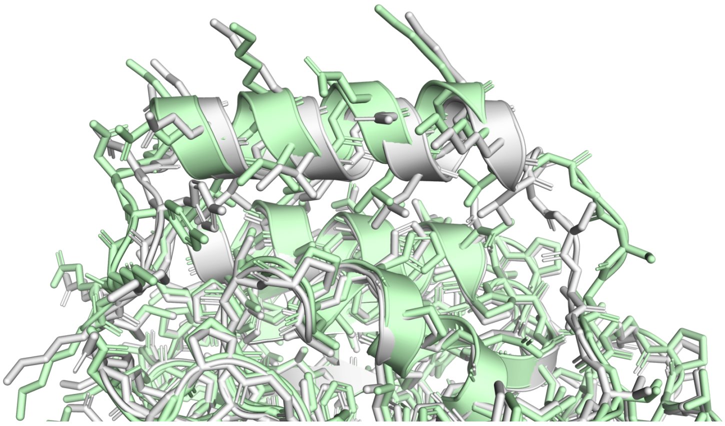

**Fig. S5. Overlay of the design model with the final circular permutation linker and the AlphaFold 3 predicted structure.**

The design model (with linker_04) is shown in grey, the AlphaFold 3 predicted structure in green (RMSD 0.530 Å).

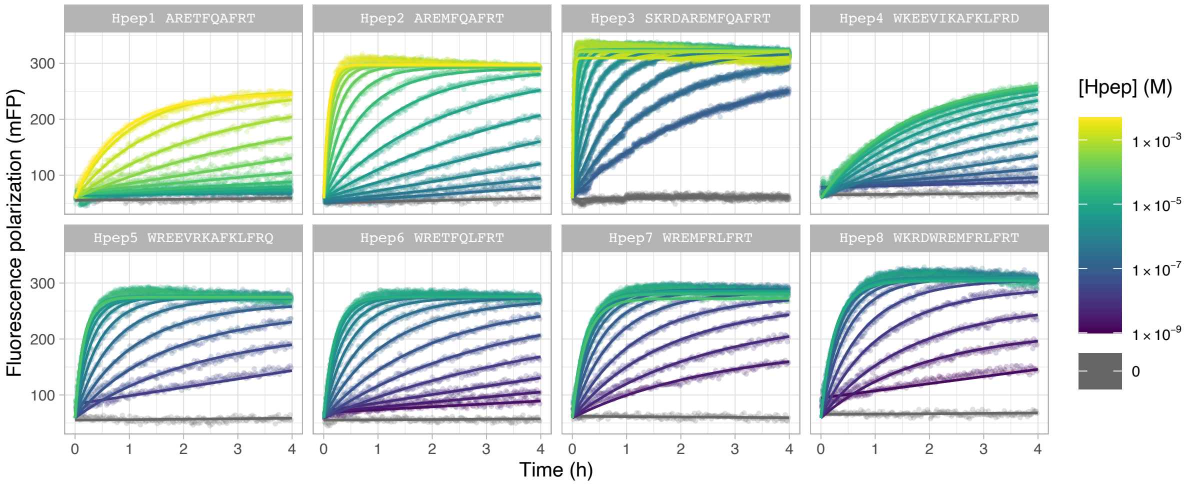


**Fig. S6. cpHaloΔ2 labeling kinetics at varying Hpep1-8 concentrations.**

Fluorescence polarization labeling kinetics of cpHaloΔ variants (10 nM) with TMR-CA (2 nM) at varying concentrations of Hpep1-8. Second-order reaction models or linear models (if reactions did not plateau) were fitted to the data to determine labeling rates.

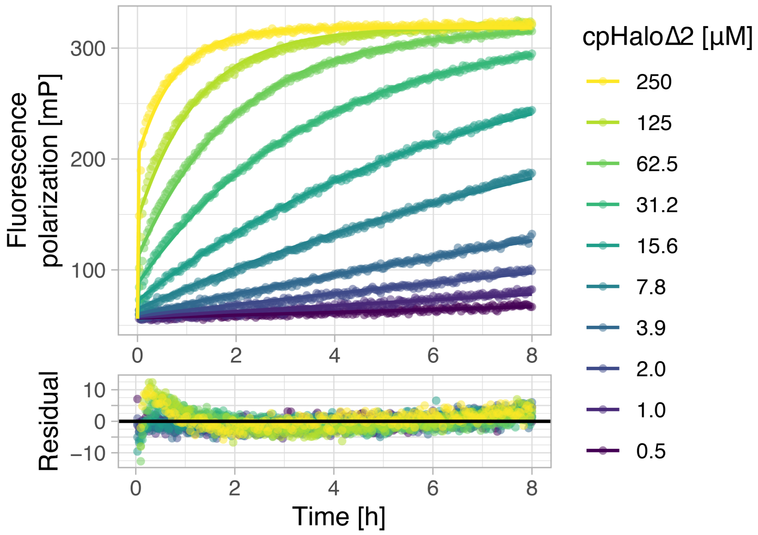


**Fig. S7. Background labeling of cpHaloΔ2 in absence of Hpep**

Fluorescence polarization labeling kinetics of cpHaloΔ2 at various concentrations with TMR-CA (50 nM). A two-step reaction model was fitted to the data globally to determine the kinetic parameters of the background labeling reaction (see table S3). Residuals of the fit are shown below.

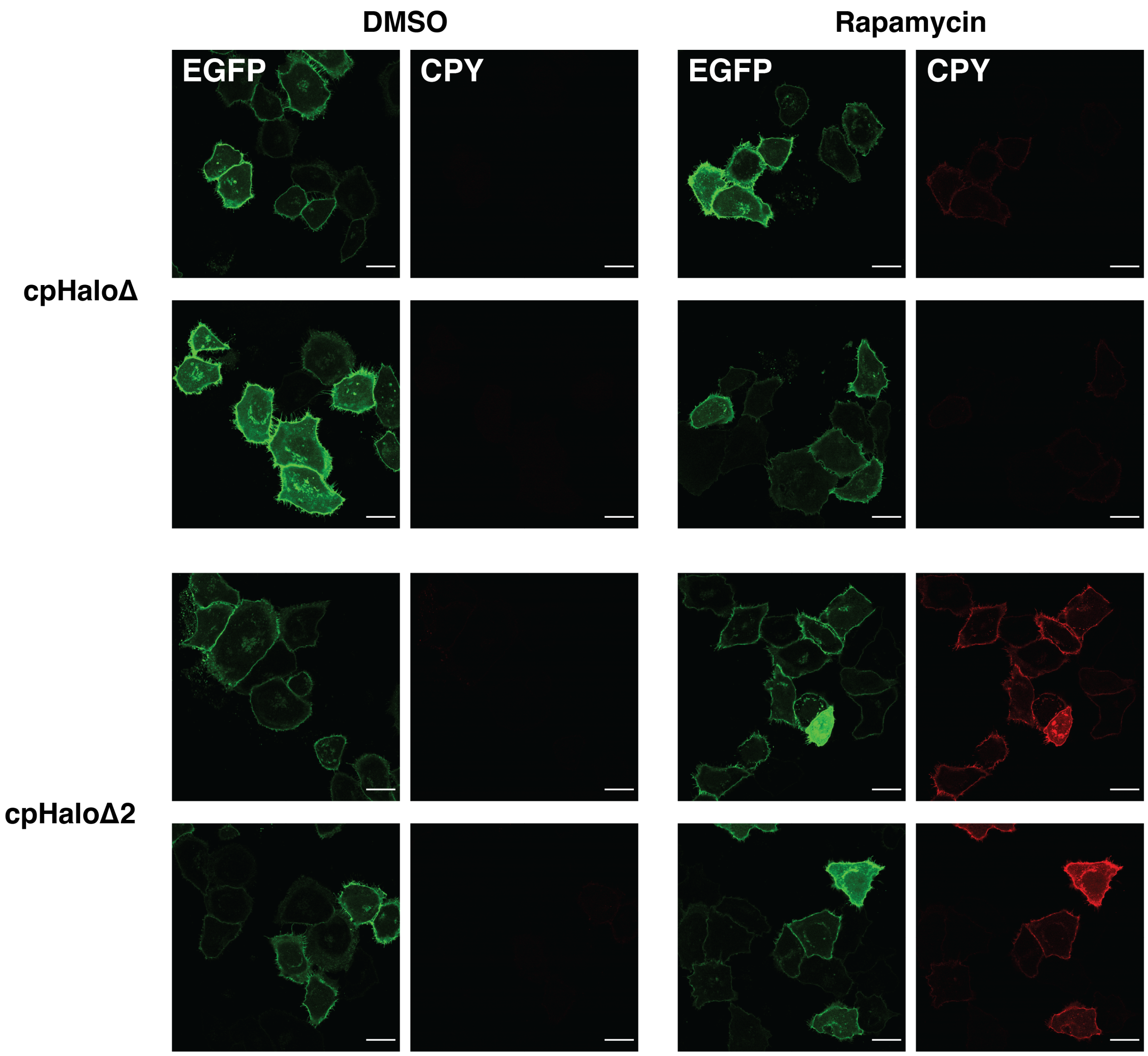


**Fig. S8. Additional fluorescence micrographs of rapamycin-dependent labeling of FKBP/FRB split-HaloTag fusions in HeLa cells.**

Additional confocal fluorescence micrographs of HeLa cells co-expressing Lyn11-EGFP-cpHaloΔ/cpHaloΔ2-$(GGS)_9$-FKBP and Hpep3-$(GGS)_3$-FRB-mScarlet. Cells were treated with CPY-CA (10 nM, 30 min) in presence or absence of rapamycin (100 nM). Scale bars are 25 µm.

**Supplementary tables**

**Table S1. Apparent second-order rate constants ($k_{app}$) of cpHaloΔ and cpHaloΔ2 at saturating concentrations of Hpep3.**

Labeling rates were determined with TMR-CA at 37 °C with a 20-minute pre-incubation before starting the reaction.

| **Variant** | **$k_{app}$** | **95% confidence interval** |
|---|---|---|
| cpHaloΔ | $0.978 \cdot 10^6$ | $(0.957 – 0.999) \cdot 10^6$ |
| cpHaloΔ2 | $1.61 \cdot 10^6$ | $(1.57 – 1.64) \cdot 10^6$ |

**Table S2. $EC_{50}$ values for the cpHaloΔ2 labeling reaction with Hpep variants 1-8.**

$EC_{50}$ values were determined with TMR-CA at 37 °C with a 20-minute pre-incubation before starting the reaction.

| Variant | Sequence | $EC_{50}$ (μM) | 95% confidence interval |
|---|---|---|---|
| Hpep1 | ARETFQAFRT | 2428 | (2077 – 2862) |
| Hpep2 | AREMFQAFRT | 283 | (262 – 306) |
| Hpep3 | SKRDAREMFQAFRT | 35.9 | (31.0 – 41.9) |
| Hpep4 | WKEEVIKAFKLFRD | 1.37 | (1.08 – 1.73) |
| Hpep5 | WREEVRKAFKLFRQ | 0.930 | (0.806 – 1.068) |
| Hpep6 | WRETFQLFRT | 0.796 | (0.710 – 0.889) |
| Hpep7 | WREMFRLFRT | 0.130 | (0.105 – 0.159) |
| Hpep8 | WKRDWREMFRLFRT | 0.0434 | (0.0383 – 0.0491) |

**Table S3. Kinetic parameters of cpHaloΔ2 background labeling in absence of Hpep.**

Parameters were determined with TMR-CA at 37 °C with a 20-minute pre-incubation before starting the reaction.

| Parameter | Value | 95% confidence interval |
|---|---|---|
| $K_D$ | 201 µM | (192 – 210) µM |
| $k_2$ | $6.55 \cdot 10^{-4}$ $s^{-1}$ | $(6.28 – 6.82) \cdot 10^{-4}$ $s^{-1}$ |
| $k_{app}$ | 3.26 $M^{-1}s^{-1}$ | (3.14 – 3.39) $M^{-1}s^{-1}$ |

**Protein sequences**

>HaloTag (as reference for numbering of mutations)

MAEIGTGFPFDPHYVEVLGERMHYVDVGPRDGTPVLFLHGNPTSSYVWRNIIPHVAPTHRCI

APDLIGMGKSDKPDLGYFFDDHVRFMDAFIEALGLEEVVLVIHDWGSALGFHWAKRNPERVK

GIAFMEFIRPIPTWDEWPEFARETFQAFRTTDVGRKLIIDQNVFIEGTLPMGVVRPLTEVEM

DHYREPFLNPVDREPLWRFPNELPIAGEPANIVALVEEYMDWLHQSPVPKLLFWGTPGVLIP

PAEAARLAKSLPNCKAVDIGPGLNLLQEDNPDLIGSEIARWLSTLEISG

>cpHaloΔ

MHHHHHHHHHHENLYFQGDVGRKLIIDQNVFIEGTLPMGVVRPLTEVEMDHYREPFLNPVDR

EPLWRFPNELPIAGEPANIVALVEEYMDWLHQSPVPKLLFWGTPGVLIPPAEAARLAKSLPN

CKAVDIGPGLNLLQEDNPDLIGSEIARWLSTLEIGGTGGSGGTGGSGGSIGTGFPFDPHYVE

VLGERMHYVDVGPRDGTPVLFLHGNPTSSYVWRNIIPHVAPTHRCIAPDLIGMGKSDKPDLG

YFFDDHVRFMDAFIEALGLEEVVLVIHDWGSALGFHWAKRNPERVKGIAFMEFIRPIPTWDE

W

His-tag, TEVp cleavage site, circular permutation linker

Designed circular permutation linkers

>linker_01

RSDDPRKTQTIASKISRDLNGS

>linker_02

KGGTKRDADKAVRDTLLSLNGQ

>linker_03

GGAPRDEALKKIEKAKRDTGDQ

>linker_04 (final linker)

KSKYDRDQILKIIAELEKKTGGS

>linker_05

KSKYDKRQIRDIADKIAKDNNHQ

>linker_06

QSKYPPEWLEKVIRELLKRKNGR

>linker_07

GADDKTKIEKILEEIKRRWQGR

>linker_08

GTSDPRNQEIAKKLARDASTVP

>linker_09

NGADKEQIDRAIEKAKRDLNNQ

>linker_10

KGASDRDEAKKLADDIRKKKGDQ

>linker_11

NSNGHRDELEKILQTIRKQNNDI

>linker_12

LKDERQRDKALEIADRADKYPTS

>cpHaloΔ2

MHHHHHHHHHHENLYFQGDVGRKLIIDQNVFIEGTLPMGVVRPLTEEEMDHYREPFLNPKDR

EPLWRFPNELPIAGEPANIVALVEEYMDWLHQSPVPKLLFWGTPGVLIPPAEAARLAKSLPN

CKAVDIGPGLNLLQEDNPDLIGSEIARWLSTLEIKSKYDRDQILKIIAELEKKTGGSIGTGF

PFDPHYVEVLGSRMHYVDVGPRDGTPVLFLHGNPTSSYVWRNIIPHVAPTHRCIAPDLIGMG

KSDKPDLGYFFDDHVRFMDAFIEALGLEEVVLVIHDWGSALGFHWAKRHPERVKGIAFMEFI

RPIPTWDEW

His-tag, TEVp cleavage site, circular permutation linker, mutations relative to cpHaloΔ

Hpep sequences

>Hpep1

ARETFQAFRT

>Hpep2

AREMFQAFRT

>Hpep3

SKRDAREMFQAFRT

>Hpep4

WKEEVIKAFKLFRD

>Hpep5

WREEVRKAFKLFRQ

>Hpep6

WRETFQLFRT

>Hpep7

WREMFRLFRT

>Hpep8

WKRDWREMFRLFRT

Lyn11-EGFP-cpHaloΔ-$(GGS)_9$-FKBP-P2A-Hpep3-$(GGS)_3$-FRB-mScarlet

MGCIKSKGKDSAGADSAGSAGMVSKGEELFTGVVPILVELDGDVNGHKFSVSGEGEGDATYG

KLTLKFICTTGKLPVPWPTLVTTLTYGVQCFSRYPDHMKQHDFFKSAMPEGYVQERTIFFKD

DGNYKTRAEVKFEGDTLVNRIELKGIDFKEDGNILGHKLEYNYNSHNVYIMADKQKNGIKVN
FKIRHNIEDGSVQLADHYQQNTPIGDGPVLLPDNHYLSTQSALSKDPNEKRDHMVLLEFVTA
AGITLGMDELYKGSGGSGDVGRKLIIDQNVFIEGTLPMGVVRPLTEVEMDHYREPFLNPVDR
EPLWRFPNELPIAGEPANIVALVEEYMDWLHQSPVPKLLFWGTPGVLIPPAEAARLAKSLPN
CKAVDIGPGLNLLQEDNPDLIGSEIARWLSTLEIGGTGGSGGTGGSGGSIGTGFPFDPHYVE
VLGERMHYVDVGPRDGTPVLFLHGNPTSSYVWRNIIPHVAPTHRCIAPDLIGMGKSDKPDLG
YFFDDHVRFMDAFIEALGLEEVVLVIHDWGSALGFHWAKRNPERVKGIAFMEFIRPIPTWDE
WGSGGTGGSGGSGGTGGSGGSGGTGGSGMGVQVETISPGDGRTFPKRGQTCVVHYTGMLEDG
KKFDSSRDRNKPFKFMLGKQEVIRGWEEGVAQMSVGQRAKLTISPDYAYGATGHPGIIPPHA
TLVFDVELLKLEGSGATNFSLLKQAGDVEENPGPGGSSKRDAREMFQAFRTGGSGGTGGSAI
LWHEMWHEGLEEASRLYFGERNVKGMFEVLEPLHAMMERGPQTLKETSFNQAYGRDLMEAQE
WCRKYMKSGNVKDLLQAWDLYYHVFRRISKGSGVSKGEAVIKEFMRFKVHMEGSMNGHEFEI
EGEGEGRPYEGTQTAKLKVTKGGPLPFSWDILSPQFMYGSRAFTKHPADIPDYYKQSFPEGF
KWERVMNFEDGGAVTVTQDTSLEDGTLIYKVKLRGTNFPPDGPVMQKKTMGWEASTERLYPE
DGVLKGDIKMALRLKDGGRYLADFKTTYKAKKPVQMPGAYNVDRKLDITSHNEDYTVVEQYE
RSEGRHSTG

Lyn11, mEGFP, cpHaloΔ, FKBP, P2A, Hep3, FRB, mScarlet

Lyn11-EGFP-cpHaloΔ2-(GGS)$_9$-FKBP-P2A-Hpep3-(GGS)$_3$-FRB-mScarlet

MGCIKSKGKDSAGADSAGSAGMVSKGEELFTGVVPILVELDGDVNGHKFSVSGEGEGDATYG
KLTLKFICTTGKLPVPWPTLVTTLTYGVQCFSRYPDHMKQHDFFKSAMPEGYVQERTIFFKD
DGNYKTRAEVKFEGDTLVNRIELKGIDFKEDGNILGHKLEYNYNSHNVYIMADKQKNGIKVN
FKIRHNIEDGSVQLADHYQQNTPIGDGPVLLPDNHYLSTQSALSKDPNEKRDHMVLLEFVTA
AGITLGMDELYKGSGGSGDVGRKLIIDQNVFIEGTLPMGVVRPLTEEEMDHYREPFLNPKDR
EPLWRFPNELPIAGEPANIVALVEEYMDWLHQSPVPKLLFWGTPGVLIPPAEAARLAKSLPN

CKAVDIGPGLNLLQEDNPDLIGSEIARWLSTLEIKSKYDRDQILKIIAELEKKTGGSIGTGF

PFDPHYVEVLGSRMHYVDVGPRDGTPVLFLHGNPTSSYVWRNIIPHVAPTHRCIAPDLIGMG

KSDKPDLGYFFDDHVRFMDAFIEALGLEEVVLVIHDWGSALGFHWAKRHPERVKGIAFMEFI

RPIPTWDEWGSGGTGGSGGSGGTGGSGGSGGTGGSGMGVQVETISPGDGRTFPKRGQTCVVH

YTGMLEDGKKFDSSRDRNKPFKFMLGKQEVIRGWEEGVAQMSVGQRAKLTISPDYAYGATGH

PGIIPPHATLVFDVELLKLEGSGATNFSLLKQAGDVEENPGPGGSSKRDAREMFQAFRTGGS

GGTGGSAILWHEMWHEGLEEASRLYFGERNVKGMFEVLEPLHAMMERGPQTLKETSFNQAYG

RDLMEAQEWCRKYMKSGNVKDLLQAWDLYYHVFRRISKGSGVSKGEAVIKEFMRFKVHMEGS

MNGHEFEIEGEGEGRPYEGTQTAKLKVTKGGPLPFSWDILSPQFMYGSRAFTKHPADIPDYY

KQSFPEGFKWERVMNFEDGGAVTVTQDTSLEDGTLIYKVKLRGTNFPPDGPVMQKKTMGWEA

STERLYPEDGVLKGDIKMALRLKDGGRYLADFKTTYKAKKPVQMPGAYNVDRKLDITSHNED

YTVVEQYERSEGRHSTG

Lyn11, mEGFP, cpHaloΔ, FKBP, P2A, Hep3, FRB, mScarlet
